# Structure of voltage-modulated sodium-selective NALCN-FAM155A channel complex

**DOI:** 10.1101/2020.07.26.221747

**Authors:** Yunlu Kang, Jing-Xiang Wu, Lei Chen

## Abstract

Resting membrane potential determines the excitability of the cell and is essential for the cellular electrical activities. NALCN channel mediates sodium leak currents, which positively tune the resting membrane potential towards depolarization. NALCN channel is involved in many important neurological processes and is implicated in a spectrum of neurodevelopmental diseases. Despite its functional importance, the mechanisms of ion permeation and voltage-modulation for NALCN channel remain elusive. Here, we report the cryo-EM structure of rat NALCN and mouse FAM155A complex to 2.7 Å resolution. The structure reveals detailed interactions between NALCN and extracellular cysteine-rich domain of FAM155A. The non-canonical architecture of NALCN selectivity filter dictates its sodium selectivity and calcium block. The asymmetric arrangement of two functional voltage-sensors confers the modulation by membrane potential. Moreover, mutations found in human diseases were mapped to the domain-domain interfaces or the pore domain of NALCN, intuitively suggesting their pathological mechanisms.

## Introduction

NALCN channel is a voltage-modulated sodium-selective ion channel (Chua et al., 2020). It mediates sodium leak currents that are essential for setting the membrane excitability (Ren, 2011). NALCN channel is involved in several crucial physiological processes including maintaining normal circadian rhythms (Nash et al., 2002) and respiratory rhythm (Lu et al., 2007). NALCN channel functions as a multi-protein complex which consists of NALCN, FAM155, UNC-79, UNC80 and other proteins (Cochet-Bissuel et al., 2014). Mutations in the genes of human NALCN or UNC-80 lead to a spectrum of neurological diseases (Bramswig et al., 2018), including infantile hypotonia, psychomotor retardation, and characteristic facies (IHPRF) (Al-Sayed et al., 2013; Koroglu et al., 2013) and congenital contractures of the limbs and face, hypotonia, and developmental delay (CLIFAHDD) (Chong et al., 2015). In addition, a gain-of function mutation of NALCN in mouse causes the *dreamless* phenotype, suggesting the function of NALCN in rapid eye movement sleep (Funato et al., 2016).

Sequence analysis shows NALCN protein shares similar domain arrangement to the pore subunit of eukaryotic voltage-gated sodium channels Na_V_ or calcium channels Ca_V_. Four consecutive voltage sensor-pore modules are fused into a single polypepetide chain. However, unique to the NALCN channel, the positive charges on voltage sensors of NALCN are much fewer than that of Na_V_ or Ca_V_. Moreover, residues in the selectivity filter of NALCN are “EEKE” instead of “DEKA” of Na_V_ or “EEEE” of Ca_V_. These features render NLACN to be a special clade in the voltage-gated channel superfamily (Kasimova et al., 2016). In addition to the NALCN subunit, FAM155, UNC-79 and UNC80 subunits are indispensible for the function of NALCN channel. The co-expression of NALCN, FAM155, UNC-79 and UNC80 proteins in heterologous expression system, such as *Xenopus laevis* oocytes or HEK293 cells, is necessary and sufficient to generate robust voltage-modulated sodium-selective currents (Chua et al., 2020). FAM155 is a transmembrane protein family with a cysteine-rich domain (CRD). There are two FAM155 family members in human, namely FAM155A and FAM155B, both of which could support the currents of NALCN complex (Chua et al., 2020). FAM155 is the homologue of NLF-1 in *c.elegans* which is required for the membrane localization of NALCN (Xie et al., 2013). It is proposed that the interaction between NALCN and FAM155 is similar to the interaction between yeast calcium channel Cch1 and its regulator Mid1 (Ghezzi et al., 2014; Liebeskind et al., 2012). The other two proteins in the complex, UNC-79 and UNC-80, are large conserved proteins without any known domains but they are essential for the function of NALCN *in vivo* (Humphrey et al., 2007; Jospin et al., 2007; Yeh et al., 2008).

Despite the functional importance and unique properties of the NALCN channel, its structure remains elusive, which impeded our understanding of the mechanism. Therefore, we embarked the structural studies of the NALCN complex.

## Results

### Structure of the NALCN-FAM155A core complex

Despite extensive screening of NALCN homologues and optimization, we failed to purify the stable hetero-tetrameric complex of NALCN-FAM155-UNC79-UNC80. However, we found C-terminal GFP tagged rat NALCN and C terminal flag tagged mouse FAM155A can generate typical NALCN currents upon co-expression of mouse UNC79 and mouse UNC80 (Fig. S1a). Moreover, rat NALCN and mouse FAM155A can form a stable core complex which can be purified chromatographically (Fig. S1b, c). Because NALCN protein dictates the properties of ion permeation, divalent ion block and voltage-modulation (Chua et al., 2020) and FAM155 is required for correct trafficking of NALCN (Xie et al., 2013), we reasoned the structure of NALCN-FAM155A complex would provide clues of how NALCN channel works. Therefore, we started with the cryo-EM studies of NALCN-FAM155 core complex. We prepared the NALCN-FAM155A complex in a buffer with sodium chloride and GDN detergent, mimicking the symmetric sodium solution used in electrophysiology recording (Fig. S1a).

Single particle cryo-EM analysis yielded a map at a nominal resolution of 2.7 Å (Figs. S2–4). The map quality was sufficient for modeling the stable portions of NALCN and FAM155A which encompass 72% of NLACN and 36% of FAM155A.

NALCN subunit shows a typical eukaryotic Na_V_ or Ca_V_ structure (Fig. 1a-d). Pore domains of domain I, II, III and IV shape the ion permeation pathway (Fig. 1a-d). Voltage sensors (VS, S1-S4 segments) pack around the pore in a domain-swapped fashion (Fig. 1a-d). Interestingly, the arrangements of VS are highly asymmetric, with voltage sensor of domain I (DI-VS) being the most off-axis (Fig. 1d). On the intracellular side, the C-terminal domain of NALCN binds onto the intracellular III-IV linker. The long II-III loop and C terminal 167 residues of NALCN were disordered. On the extracellular side, the CRD of FAM155A protrudes from the top of the complex into the extracellular space while the two predicted transmembrane helices of FAM155A were invisible, probably due to their flexibility.

**Fig. 1.**
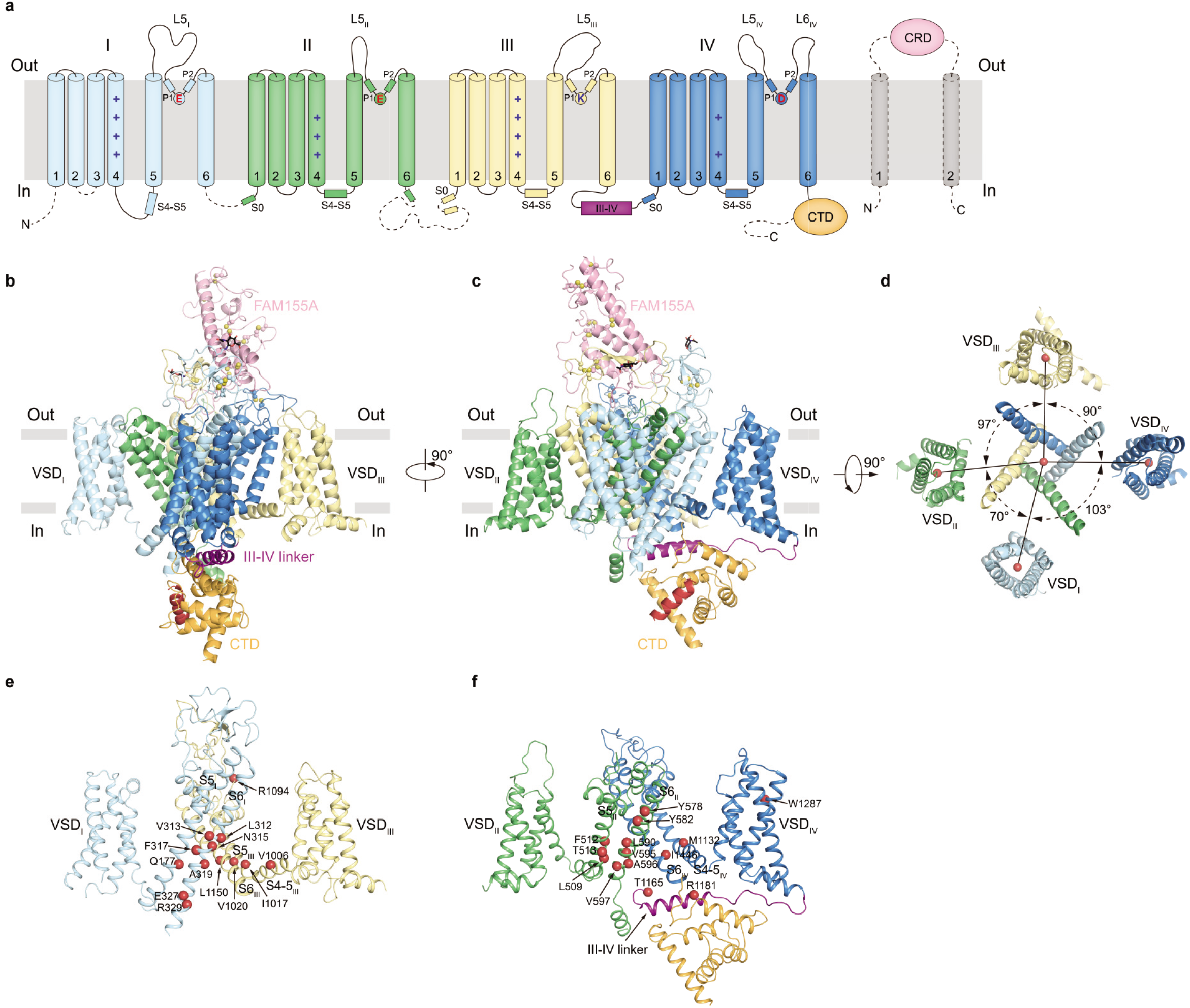
Overall structure of NALCN and FAM155A complex. **a**, Topology of NALCN and FAM155A subunits. Helices are shown as cylinders, unmodeled disordered regions are shown as dashed lines. The phospholipid bilayer is shown as grey layers. CTD, C-terminal intracellular domain of NALCN. CRD, cysteine rich domain of FAM155A. Plus signs represent positively charged residues on S4 segments. Key residues on selectivity filter are indicated. Dashed lines indicate flexible regions with invisible densities in our cryo-EM map. **b**, Side view of NALCN and FAM155A complex. Sugar moieties are shown as black sticks, disulfide bonds are shown as golden spheres. The approximate boundaries of phospholipid bilayer are indicated as grey thick lines. **c**, A 90°rotated view compared to **b**. **d**, The arrangement of NALCN transmembrane domain illustrated in the top view. For clarity, only four voltage sensing domains and S6 segments are shown. The angles between adjacent voltage sensing domains were labeled. Angular measurements were based on the center of mass positions (red spheres) of each domain. **e**, **f**, Structural mapping of disease-related mutations in NALCN. For clarity, only two nonadjacent domains are shown in each panel. The Cα atoms of disease-related residues are shown as red spheres.

The structure of NALCN allowed us to locate all of the reported missense mutations identified in human diseases and model animals (Fig. 1e, f) (Al-Sayed et al., 2013; Aoyagi et al., 2015; Campbell et al., 2018; Chong et al., 2015; Fukai et al., 2016; Funato et al., 2016; Karakaya et al., 2016; Lozic et al., 2016; Sivaraman et al., 2016; Vivero et al., 2017; Wang et al., 2016; Yeh et al., 2008).

### Structure of FAM155A CRD and its interaction with NALCN

The FAM155A has an extracellular cysteine-rich domain (CRD) which is folded by three visible consecutive α helices and the connecting loops in between. The CRD of FAM155A is highly cross-linked by disulfide bonds owing to the high content of cysteines in this domain. Most of the Cys residue pairs are conserved from NLF-1 of C.elegans to FAM155A/B in human, suggesting their important functions, presumably by stabilizing the structure of CRD (Fig. 2a-c, Fig. S6).

**Fig. 2.**
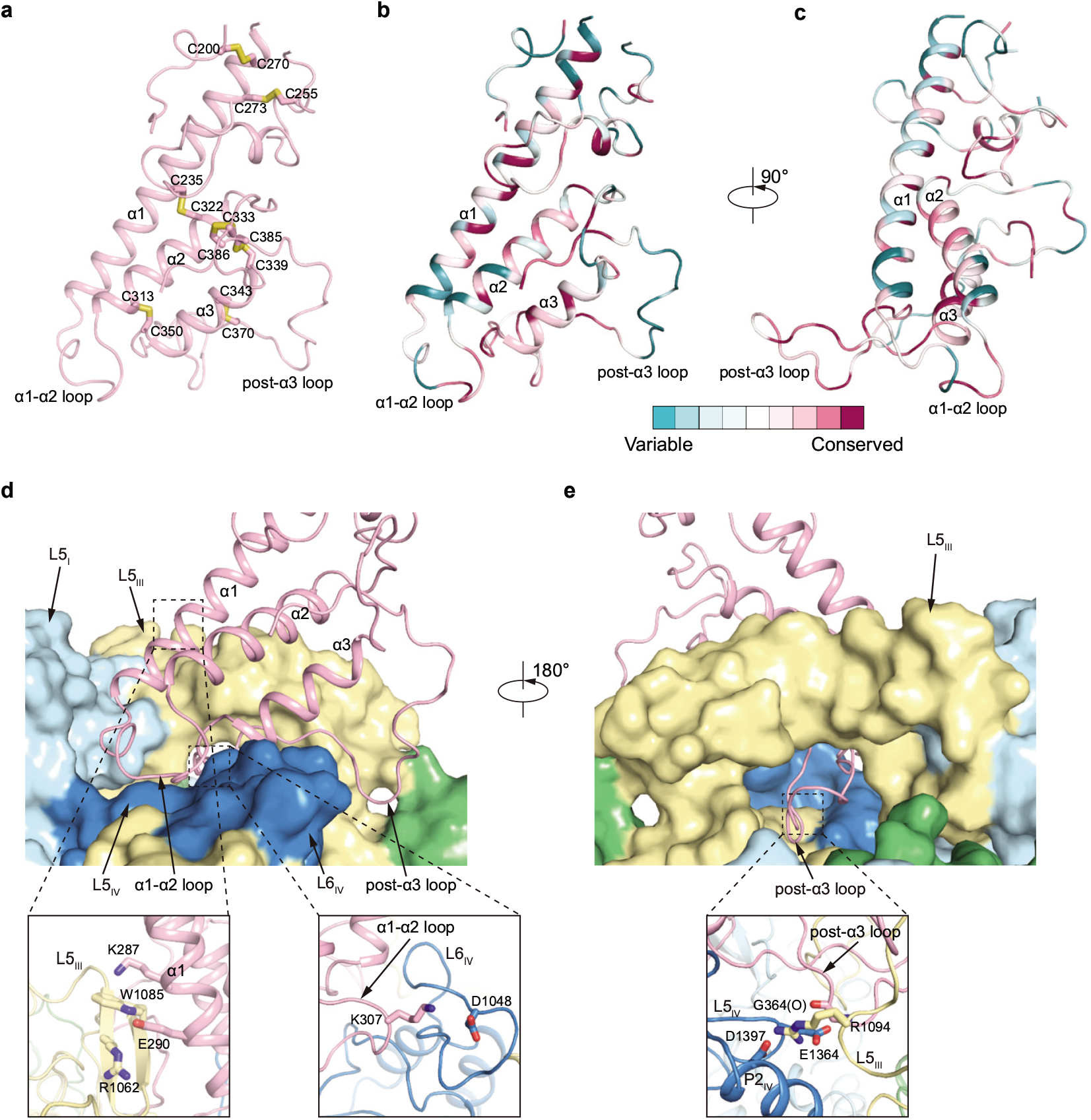
Structure of FAM155A CRD and its interactions with NALCN. **a**, Structure of FAM155A CRD (residues 190-201, 227-260 and 267-389). Disulfide bonds are shown as yellow sticks. **b**, Structural mapping of conserved residues of FAM155 family members, generated by ConSurf (Ashkenazy et al., 2010). **c**, A 90°rotated view compared to **b**. **d**, Interactions between FAM155A and NALCN. FAM155A is shown as cartoon. NALCN is shown as surface. Key interaction regions are boxed and zoomed in the close-up views below. **e**, A 180°rotated view compared to **d**.

The CRD of FAM155A binds at the top of NALCN channel (Fig. 1b, c). Extracellular loops of domain I, III and IV (L5_I_, L5_III_, L5_IV_, and L6_IV_) in NALCN form a platform for FAM155A binding (Fig. 2d). The post-α3 loop of FAM155A extends over the extracellular ion entrance of the NALCN pore (Fig. 2e). The L5_III_ of NALCN covers above the post-α3 loop of FAM155A and locks it in the sandwiched conformation (Fig. 2e). The binding of CRD of FAM155A onto NALCN relies on its extensive interactions with L5_III_, L5_IV_ and L6_IV_ of NALCN. In detail, K287 and E290 on α1 of FAM155A make cation-π and electrostatic interactions with W1085 and R1062 on L5_III_, respectively (Fig. 2d). K307 on α1-α2 loop interacts with D1408 on L6_IV_ (Fig. 2d). R1094 on L5_III_ interacts with the carbonyl group of G364 of FAM155A, D1397 on P2_IV_ and E1364 on L5_IV_ (Fig. 2e). These interacting residues are highly conserved (Fig. 2b-c, Fig. S5–6). Notably, it is reported that mutation of R1094Q, which locates on the interface between FAM155A, domain III and domain IV (Fig. 2e), leads to disordered respiratory rhythm with central apnea (Campbell et al., 2018), suggesting the importance of these inter-subunit interactions.

### The pore domain of NALCN

Along the ion permeation pathway, negatively charged residues are broadly distributed. They may function to attract sodium ions for permeation (Fig. 3a). The calculated pore profile reveals two constrictions (Fig. 3b-c). The constriction close to the extracellular side is the selectivity filter. At this layer, the side chains of E280 (domain I), E554 (domain II), K1115 (domain III) and D1390 (domain IV) point to the center of the pore (Fig. 3c-f). In contrast, E1389 of domain IV has two possible alternative conformations, either pointing up, making electrostatic interaction with R1384 on P1 helix of domain IV, or pointing down (Fig. 3c-f), none of which points to the center of pore. This is distinct from previous prediction based on sequence alignments and structures of Na_V_ and Ca_V_, which suggested that E1389 on domain IV is part of the selectivity filter (Fig. 3d). The conformation and structural arrangement of E1389 and D1390 in NALCN was not observed previously in other Na_V_ or Ca_V_ channels. It is recently reported that the sodium current of NALCN can be blocked by extracellular calcium at physiological concentration and mutations of D1390A has more profound effect than D1389A (Chua et al., 2020), which is in agreement with our observations showing D1390 contributes to ion permeation.

**Fig. 3.**
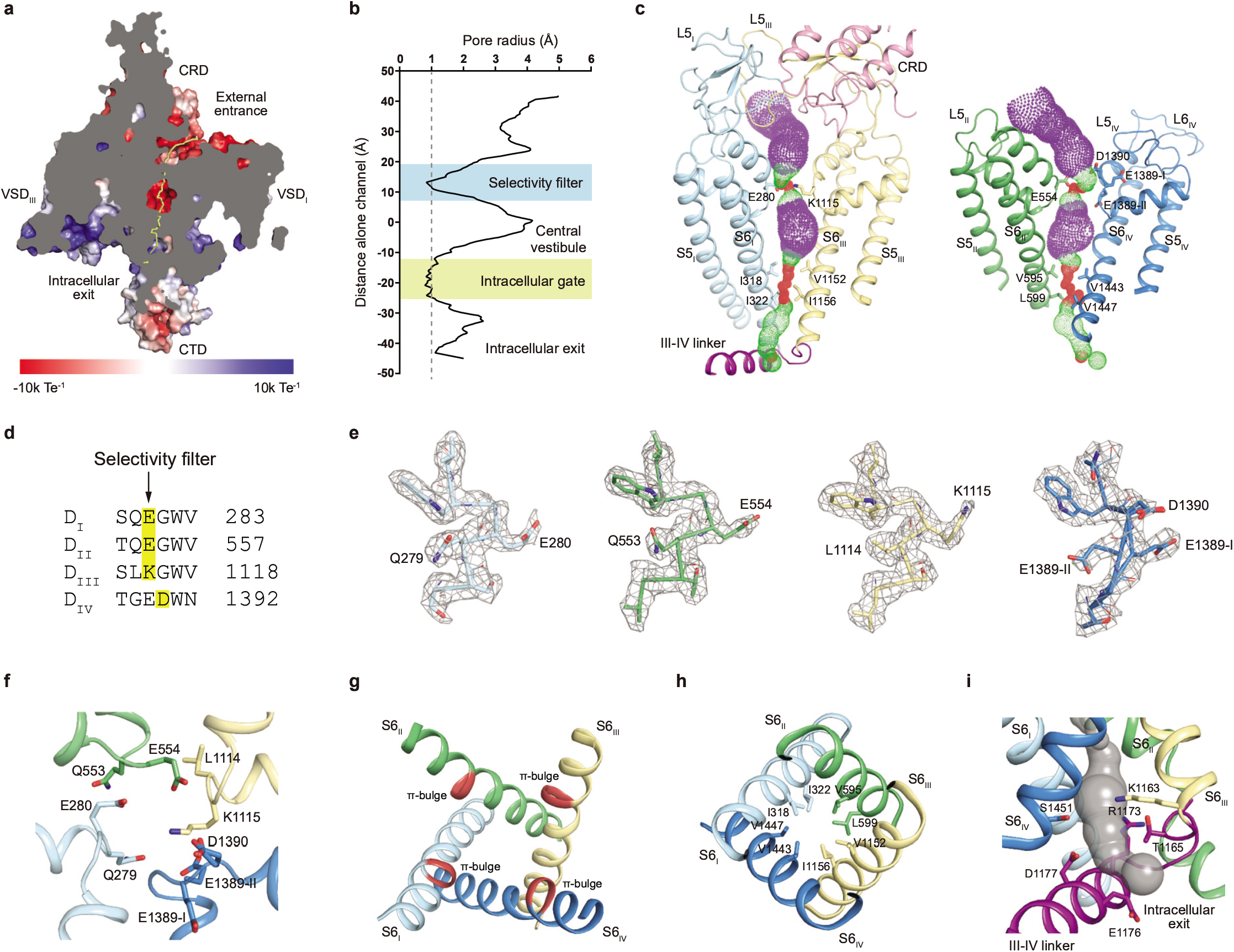
Structure of the pore domain of NALCN. **a**, A cut-open side view of NACLN and FAM155A complex. The surface electrostatic potential is calculated by APBS (Jurrus et al., 2018). The ion permeation pathway is shown as yellow line. **b**, Pore profile of NALCN channel shows two constrictions. **c**, The ion permeation pathway of the pore domain, calculated with HOLE (Smart et al., 1993), is shown as color dots. Purple, green and red dots represent pore radii greater than 2.8 Å, 1.4 to 2.8 Å and less than 1.4 Å, respectively. **d**, Sequence alignment of the residues around the selectivity filter of rat NALCN. Residues that participate in ion permeation is highlighted in yellow. Predicted residues for selectivity filter are indicated with arrow. **e**, Densities of selectivity filter of each repeat, shown as grey mesh, are contoured at 3.5 σ. **f**, Top view of selectivity filter. Key residues are shown as sticks. **g**, Structure of S6 segments. π-bulges are highlighted in red. **h**, Top view of the intracellular gate, hydrophobic residues forming the narrowest constrictions are shown as sticks. **i**, Side view of the intracellular exit for sodium. Grey surface represents the calculated ion permeation pathway.

At the level of central vestibule, the S6 helices of all of the four domains show π-bulges in the middle (Fig. 3g). It is suggested that the α-π helix transition is important for the gating of some channels (Zubcevic and Lee, 2019), but its function in NALCN remains unclear. The constriction close to the cytoplasmic side is at the level of bundle crossings which represents the putative intracellular gate of NALCN. The gate is tightly sealed by several hydrophobic residues on S6 helices (Fig. 3b, c, h), indicating the current structure represents a non-conductive state. Below the gate, the intracellular exit of the ion permeation pathway is blocked by III-IV linker at one side but remains laterally open towards the direction of domain III (Fig. 3i). We found many disease mutations of NALCN accumulate in the pore, around the bundle crossing region, suggesting they might exert their pathological functions by modulate NALCN gating directly (Fig. 1e, f).

### Conformations of degenerated voltage sensors at depolarizing membrane potential

VS senses changes of membrane potential and drive the gating movement of voltage-gated ion channels. Previous studies showed that the consecutive positive gating charges reside on one face of the S4 helix. On the opposing S2 helix, negative or polar residues are separated by an aromatic residue (Phe or Tyr) and followed by a positively charged residue, forming the E1-X2-F3-E4-R5 structural motif. Together with a negatively charged residue on S3 helix, F3 and E4 on S2 form the so-called charge transfer center (CTC) at the bottom of VS (Tao et al., 2010). The S4 helix of functional VS is in the “down” conformation due to the negative voltage at resting membrane potential, while at depolarizing potential, S4 helix moves “up” in response to the changes of electrostatic field (Lee and MacKinnon, 2019; Wisedchaisri et al., 2019). In contrast to the canonical voltage-gated ion channels, some of the voltage sensors of NALCN are degenerated due to mutations of key residues in either charge-transfer center on S2 or gating charges on S4. However, NALCN is modulated by voltage (Chua et al., 2020). Therefore, the structure of NALCN provides a unique opportunity to explore the conformations of degenerated VS at 0 mV, a depolarizing state.

Voltage sensor of domain I (DI-VS) has a consensus S2 helix. On S4 helix, positively charged residues of R2, R3 and R5 are conserved, while the middle R4 is replaced by Ile (I149) (Fig. 4a). We observed R2 (R143) interacts with E128 on S3, R3 (R148) interacts with E1 (D75) on S2 and R5 (R152) interacts with E4 (E85) on S2. The Cα of F3 (Y82) is at the level similar to the Cα of R4 (I149) (Fig. 4b). For reference, the Cα of F3 (Y168) in DI-VS of Na_V_1.4 in depolarizing state (PDB ID:6AGF) is at a similar level to R3 (R225), suggesting S4 of NALCN DI-VS is even “upper” than Na_V_1.4 (Fig. S7a). The conserved structural features and depolarized conformation of DI-VS suggests it is a functional voltage sensor. Indeed, recent studies showed wild type NALCN responses to voltage pulse at holding potential of 0 mV but the R146Q+R152Q (R3+R5) mutant of DI-VS does not (Chua et al., 2020). The S4-S5 linker of available Na_V_ or Ca_V_ structures adopts a helical structure which is parallel to the membrane plane. However, the N terminus of S4-S5 linker of NALCN DI shows a loop-like structure while its C terminus is fused with S5 to form an exceptionally long S5 helix (Fig. S7b). Together with the notion that DI-VS is off-axis (Fig. 1d), these structural observations indicate DI-VS of NLACN might convey the voltage signal to the pore in a manner that is distinct from other VSs.

**Fig. 4.**
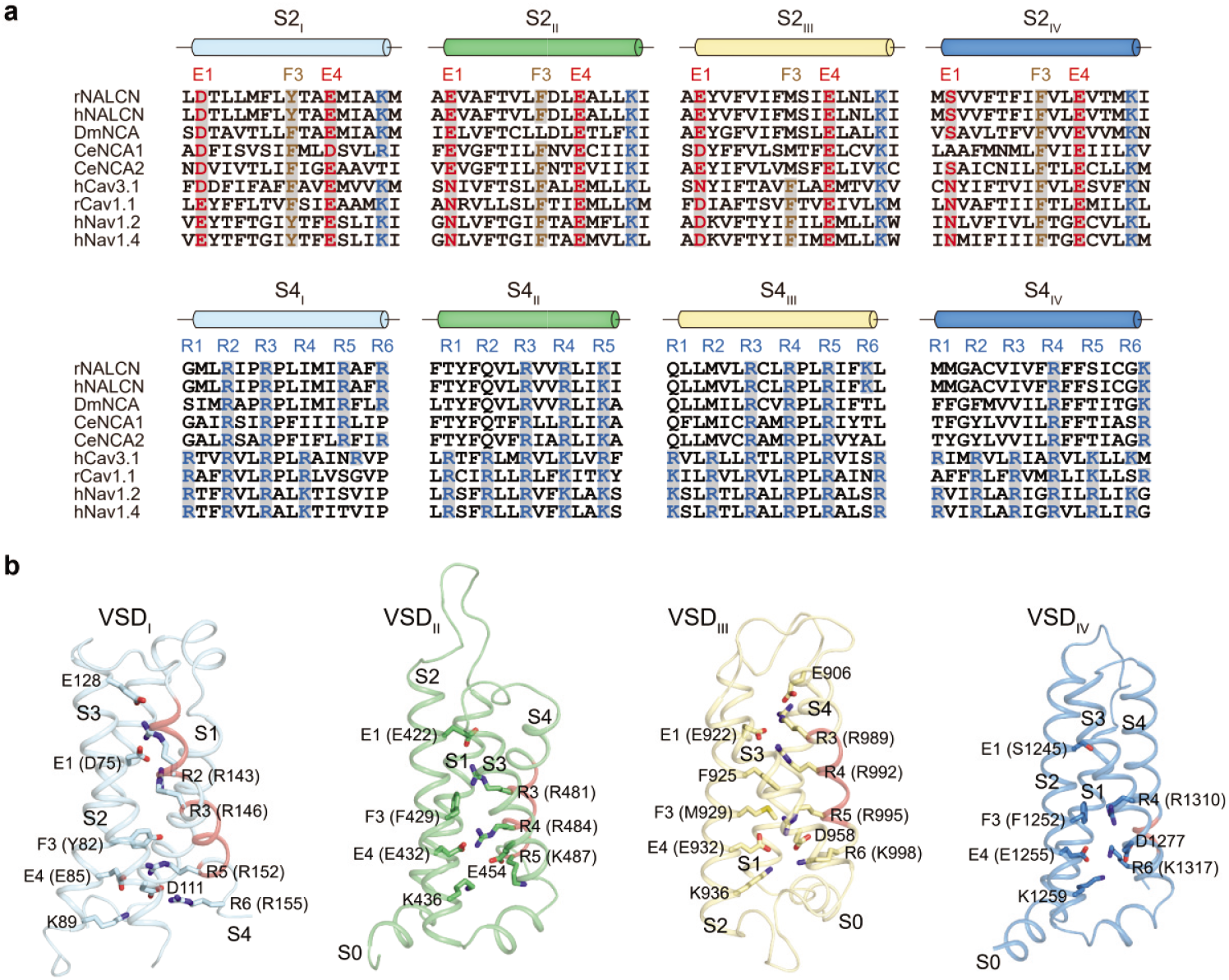
Voltage-sensors of NALCN at the depolarized state. **a**, Sequence alignment of S2 and S4 segments. The sequences of rat NALCN, human NALCN, fruit fly NCA, nematode NCA1, NCA2, human Ca_v_3.1, rabbit Ca_v_1.1, human Na_v_1.2 and human Na_v_1.4 were aligned using PROMALS3D (Pei and Grishin, 2014). Residues forming the charge transfer center are shaded in grey. The conserved negative or polar E1 and E4 on S2 segments, F3 on S2 segments and positive R1-R6 on S4 segments are highlighted in red, brown and blue, respectively. **b**, Structures of four VSDs in NALCN. Gating charges and charge transfer center residues are shown as sticks. 3_10_ helices are colored in pink.

DII-VS has an intact CTC and three consecutive gating charges on S4 (R3-R5) and the Cα of F3 (F429) is at similar level to Cα of R3 (R481), which is similar to DII-VS of Na_V_1.4 (PDB ID: 6AGF) (Fig. 4, Fig. S7a). These observations suggest DII-VS is a functional VS. Correlating with this, functional studies showed mutation of R3 (R481) together with one of R4 (R484) or R5 (K487) render NALCN unresponsive to pulse at 0 mV(Chua et al., 2020).

Although DIII-VS of NALCN has three conserved consecutive gating charges on S4 (R3-R5), it has a defective CTC on S2 and the essential aromatic residue at F3 is replaced by M929 (Fig. 4a). This is consistent with functional studies showing neutralizing mutation of all the gating charges on S4 (R989Q+R992Q+R995Q) has little effect on the voltage modulation of NALCN channel (Chua et al., 2020). We observed S4 residues above R3 (R989) adopt α helical structure while residues below R3 form a 3_10_ helix. R3 interacts with E906 on S1-S2 linker and E1 (E922). R4 interacts with E1 (E922) and F925. R5 is stabilized by interactions with E4 and D958 on S3. The Cα of F3 (M929) is at the same level to Cα of R5 (R995), which is similar to Na_V_1.4 (PDB ID: 6AGF) (Fig. 4b, Fig. S7a), suggesting DIII-VS of NALCN is in an “up” conformation, although its CTC is defective.

In contrast to the DIII-VS which has a defective CTC but three conserved gating charges, the DIV-VS of NLACN has conserved CTC but only two gating charges R4 (R1310) and R6 (K1317) on S4. Moreover, the majority of S4 adopts an α helical structure on which the spatial arrangement of gating charges is not optimal for movements upon membrane potential changes, indicating DIV-VS is defective. In agreement with this, mutation of R1310Q has no effect on the voltage sensitivity of NALCN (Chua et al., 2020). Notably, it is reported the W1287L mutation on S3 of DIV-VS can leads to the loss of currents and IHRRF (Al-Sayed et al., 2013; Bouasse et al., 2019) (Fig. 1f), probably by affecting protein folding or stability.

### Intracellular domains

At the intracellular side of domain III, the well-ordered III-IV linker makes two sharp turns at G1164 and I1168 and reroutes the intracellular exit for sodium towards the direction of domain III (Fig. 3c). There is an extensive polar interaction network between III-IV linker and the intracellular side of domain I and II, involving E327 and R329 of domain I, E603, K609 and Q613 of domain II, Q1172 and R1173 of III-IV linker. Residues following this turn of III-IV linker fold into the III-IV helix which binds below domain IV. R1174 on this helix interacts with E1458 on CTD while R1181 interacts with main chain carbonyl group of S1451, L1452 and Y1454 (Fig. 5a). The gain-of-function R1181Q mutation on this helix causes intellectual disability, episodic and persistent ataxia, arthrogryposis, and hypotonia (Aoyagi et al., 2015).

**Fig. 5.**
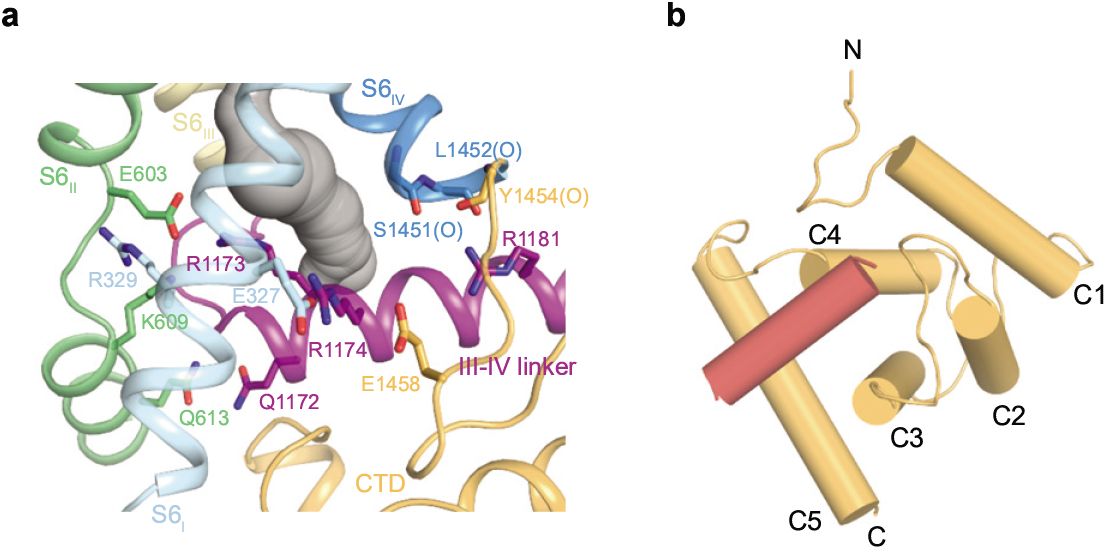
Structure of NALCN intracellular domains. **a**, III-IV linker has extensive polar interactions with CTD and S6 segments in NALCN. The interacting residues are shown as sticks. **b**, Structure of CTD of NALCN. Helices are shown as cylinders.

Moreover, T1165P mutation can lead to CLIFAHDD syndrome, further emphasizing the important function of III-IV linker. The position of III-IV linker in NALCN shares similarity to Ca_V_1.1 (PDB: 5GJV)(Wu et al., 2016) and Na_V_PaS (PDB: 5X0M)(Shen et al., 2017) channels but distinct from Na_V_1.2 (PDB: 6J8E)(Pan et al., 2019), Na_V_1.4 (PDB: 6AGF)(Pan et al., 2018), Na_V_1.5 (PDB: 6UZ3)(Jiang et al., 2020) and Na_V_1.7 (PDB: 6J8G)(Shen et al., 2019) (Fig. S8a). NALCN does not have the IFM motif on the III-IV linker, which is involved in fast inactivation of Na_V_. This is in accordance with the fact that NALCN does not show fast inactivation during depolarization.

The III-IV helix, together with the intracellular surface of NALCN form a platform for the binding of CTD. The resolved portion of NALCN CTD forms a calmodulin-like helical domain that packs below III-IV linker (Fig. 5b). The CTD shares similarity to that of Ca_V_1.1 (PDB ID: 5GJV) (Wu et al., 2016) and Na_V_PaS (PDB ID: 5X0M) (Shen et al., 2017), except the C terminal helix orients differently.

## Discussion

The high-resolution structure of NALCN-FAM155A complex presented here shows the architecture of the NALCN channel core components, reveals the structure of FAM155 CRD and depicts the detailed inter-molecular interface between NALCN and FAM155A subunits. More importantly, the non-canonical configuration of selectivity filter suggests the mechanism of sodium permeation and calcium block. Furthermore, the asymmetric spacial arrangement of both functional and degenerate voltage sensors provides mechanistic clues for the voltage-sensitivity of NALCN. In addition, our structure also provides a template for drug discovery targeting NALCN channel in related human diseases.

## Methods

### Cell culture

Sf9 insect cells (Thermo Fisher Scientific) were cultured in SIM SF (Sino Biological) at 27°C. HEK293F suspension cells (Thermo Fisher Scientific) were cultured in Freestyle 293 medium (Thermo Fisher Scientific) supplemented with 1% FBS at 37°C with 6% CO_2_ and 70% humidity. HEK293T (ATCC) cells were cultured in Dulbecco’s Modified Eagle Medium (DMEM, Thermo Fisher Scientific) supplemented with 10% FBS and 1% penicillin-streptomycin (Thermo Fisher Scientific) at 37°C with 5% CO_2_. The cell lines were routinely checked to be negative for mycoplasma contamination.

### Electrophysiology

HEK293T cells were co-transfected with rNALCN-GFP, mFAM155A-FLAG, mUNC80 and HA-mUNC79 plasmids at a ratio of 1:1:1:1 using polyethylenimine (PEI 25K, Polysciences) and incubated for 24h before recording. Patch electrodes were pulled with a horizontal microelectrode puller (P-1000, Sutter Instrument Co, USA) to a resistance of 3-5 MΩ. Wholecell patch clamps were performed using an Axon-patch 200B amplifier (Axon Instruments, USA), and data were collected with pClamp 10 software (Axon Instruments, USA) and an Axon Digidata 1550B digitizer (Axon Instruments, USA). Pipette solution containing (mM): 10 HEPES (pH 7.2, NaOH), 136 NaCl, 10 NaF, 5 EGTA, 2 Na_2_ATP and bath solution containing (mM): 10 HEPES (pH 7.4, NaOH), 150 NaCl, 30 glucose. In some experiments, 10 μM GdCl_3_ (Sigma) was added to the bath solution to block the inward current. Signals were acquired at 5 kHz and low-pass filtered at 500 Hz.

### Protein expression and purification

Mouse FAM155A was cloned from the cDNA that was reverse transcribed from the total mRNA of mouse brain. The cDNAs of rat NALCN and mouse FAM155A were cloned into a modified BacMam expression vector with C-terminal GFP-strep and FLAG tag, respectively (Goehring et al., 2014; Li et al., 2017). The baculoviruses were produced using the Bac-to-Bac system and amplified in Sf9 cells. For protein expression, HEK293F cells at density of 2.5 × 10^6^ ml^-1^ were infected with 6% volume of NALCN P2 virus and 4% volume of FAM155A P2 virus. 10 mM sodium butyrate was added to the culture 12 hours post infection and transferred to 30 °C incubator for another 48 hours before harvesting. Cells were collected by centrifugation at 4,000 rpm (JLA-8.1000, Beckman) for 10 min, and washed with 50 mM Tris (pH 7.5), 150 mM NaCl, 2 μg/ml aprotinin, 2 μg/ml pepstatin, 2 μg/ml leupeptin, flash frozen and storage at −80°C.

For each batch of protein purification, cell pellet corresponding to 0.75 liter culture was thawed and extracted with 54 ml lysis buffer containing 50 mM Tris (pH 7.5), 150 mM NaCl, 2 μg/ml aprotinin, 2 μg/ml pepstatin, 2 μg/ml leupeptin, 10% (v/v) glycerol, 1 mM phenylmethanesulfonyl fluoride (PMSF) and 1% (w/v) glyco-diosgenin (GDN, Anatrace) at 4 °C for 80 min. 1 mM iodoacetamide was added during the detergent extraction procedure to reduce non-specific cysteine crosslinking. Then the cell debris was removed by centrifugation at 16,000 rpm (JA-25.50, Beckman) for 10 min. The supernatant was ultra-centrifuged at 35,000 rpm (Type 45 Ti, Beckman) for 40 min. The solubilized proteins were loaded onto 3ml Streptactin Beads 4FF (Smart-Lifesciences) column and washed with 15 ml W buffer, which containing 20 mM Tris (pH 7.5), 150 mM NaCl, 2 μg/ml aprotinin, 2 μg/ml pepstatin, 2 μg/ml leupeptin, 10% glycerol and 0.02% GDN. The column was washed with 50 ml W buffer plus 10 mM MgCl_2_ and 2 mM adenosine triphosphate (ATP) to remove contamination of heat shock proteins. Then the column was washed with 20 ml W buffer again to remove residual MgCl_2_ and ATP. The target protein was eluted with 12 ml W buffer plus 40 mM Tris (pH 8.0) and 5 mM D-desthiobiotin (IBA). Eluted protein was concentrated using 100-kDa cut off concentrator (Millipore) and further purified by Superose 6 increase (GE Healthcare) running in a buffer containing 20 mM Tris (pH 7.5), 150 mM NaCl and 0.02% GDN. Fractions corresponding to monomeric NALCN-FAM155A complex were pooled and concentrated to A_280_ = 5.

### EM sample preparation

Holey carbon grids (Quantifoil Au 300 mesh, R 0.6/1) were glow-discharged by Solarus advanced plasma system (Gatan) for 120 s using 50% O_2_ and 50% Ar. Aliquots of 3μl concentrated protein sample were applied on glow-discharged grids and the grids were blotted for 2s before plunged into liquid ethane using Vitrobot Mark IV (Thermo Fisher Scientific).

### Cryo-EM data acquisition

Cryo-grids were firstly screened on a Talos Arctica electron microscope (Thermo Fisher Scientific) operating at 200 kV with a K2 Summit direct electron camera (Thermo Fisher Scientific). The screened grids were subsequently transferred to a Titan Krios electron microscope (Thermo Fisher Scientific) operating at 300 kV with a K2 Summit direct electron camera and a GIF Quantum energy filter set to a slit width of 20 eV. Images were automatically collected using SerialEM in super-resolution mode at a nominal magnification of 130,000×, corresponding to a calibrated super-resolution pixel size of 0.5225 Å with a preset defocus range from −1.5 μm to −1.8 μm. Each image was acquired as a 7.12 s movie stack of 32 frames with a dose rate of 7.15 e^-^Å^-2^s^-1^, resulting in a total dose of about 50.9 e^-^Å^-2^.

### Image processing

The image processing workflow is illustrated in Fig. S3. A total of 5,470 super-resolution movie stacks were collected. Motion-correction, two-fold binning to a pixel size of 1.045 Å, and dose weighting were performed using MotionCor2 (Zheng et al., 2017). Contrast transfer function (CTF) parameters were estimated with Gctf (Zhang, 2016). Micrographs with ice or ethane contamination, and empty carbon were removed manually. A total of 2,226,376 particles were auto-picked using Gautomatch from 5,428 micrographs. All subsequent classification and reconstruction was performed in Relion 3.1 (Zivanov et al., 2018) unless otherwise stated. Two rounds of reference-free 2D classification were performed to remove contaminants, yielding 567,397 particles. The particles were subjected to 57 iterations K = 1 global search 3D classification with an angular sampling step of 7.5° to determine the initial alignment parameters using the initial model generated from cryoSPARC (Punjani et al., 2017). For each of the last seven iterations of the global search, a K = 6 multi-reference local angular search 3D classification was performed with an angular sampling step of 3.75° and search range of 30°. The multi-reference models were generated using reconstruction at the last iteration from global search 3D classification low-pass filtered to 8 Å, 15 Å, 25 Å, 35 Å, 45 Å and 55 Å, respectively. The classes that showed obvious secondary structure features were selected and combined. Duplicated particles were removed, yielding 292,877 particles in total. These particles were subjected to additional three rounds of multi-reference local angular search 3D classification using the same parameters. Each round had 30 iterations, and the particles corresponding to good classes from the last 15 iterations of each round were combined, and duplicated particles were removed, yielding 201,654 particles. These particles were subsequently subjected to local search 3D auto-refinement, which resulted in a map with resolution of 3.0 Å. In order to further clean up the dataset, three rounds random-phase 3D classification were performed with K = 2, an angular sampling step of 1.875° and local search range of 10°, using two reference models, which are the model obtained from previous refinement and the phase-randomized model. Phase-randomized models were generated from the model obtained from previous refinement using randomize software (from the lab of Nikolaus Grigorieff) for phase-randomization beyond 40 Å, 30 Å and 20 Å for the first, second and third round of 3D classification. For random-phase 3D classification, each round had 30 iterations, and the particles corresponding to good class from the last 15 iterations of each round were combined, and duplicated particles were removed, yielding 135,043 particles, resulting in a 2.9 Å map. CTF refinement was then performed with Relion 3.1, the resolution was improved to 2.8 Å. The particles were then re-extracted, recentered, and re-boxed from 240 pixels to 320 pixels. After 3D auto-refinement, the resolution reached 2.7 Å.

All of the resolution estimations were based on a Fourier shell correlation (FSC) of 0.143 cutoff after correction of the masking effect. B-factor used for map sharpening was automatically determined by the post-processing procedure in Relion 3.1 (Zivanov et al., 2018). The local resolution was estimated with Relion 3.1 (Zivanov et al., 2018).

### Model building

The initial model of NALCN alone was generated by SWISS-MODEL based on the structure of rabbit Ca_v_1.1-α1 subunit (PDB ID: 5GJV) (Wu et al., 2016). The model was then fitted into the cryo-EM map using Chimera (Pettersen et al., 2004) and rebuilt manually using Coot (Emsley et al., 2010). Model of FAM155A was build manually using Coot (Emsley et al., 2010). Model refinement was performed using phenix.real_space_refine in PHENIX (Adams et al., 2010).

---

**Fig. S1.**
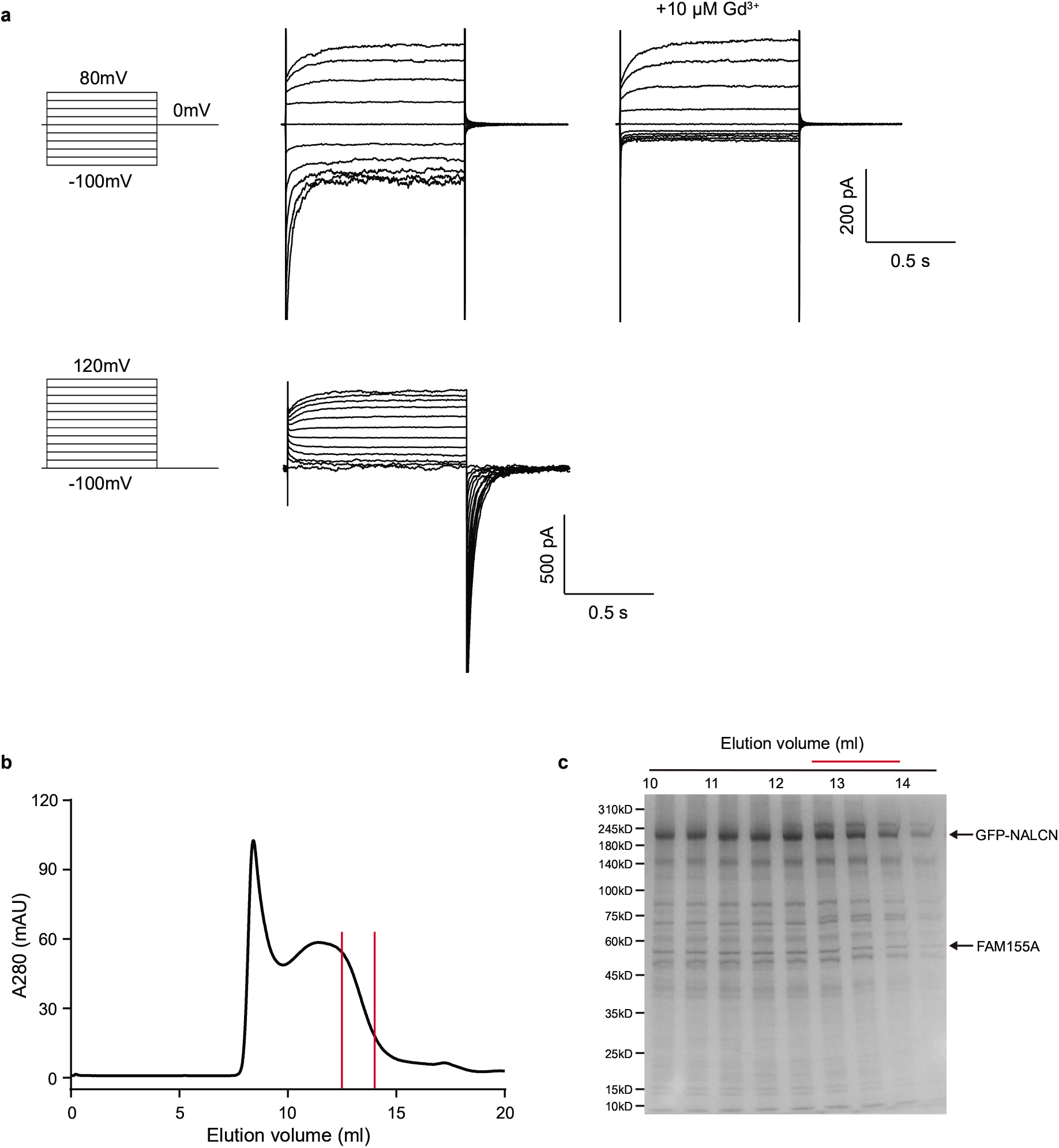
Electrophysiology and biochemistry characterization of NALCN channel complex. **a**, Representative current traces of HEK293 cells expressing C-terminal GFP-tagged rNALCN, C-terminal FLAG-tagged mFAM155A, mUNC80 and N-terminal HA-tagged mUNC80 in the symmetric sodium solutions. 10 μM GdCl_3_ were added to the bath solution to block the inward current. **b**, Size-exclusion chromatography of NALCN-FAM155A complex on a Superose 6 increase column. The fractions between the red vertical lines were pooled and concentrated for cryo-EM sample preparation. **c**, SDS-PAGE of fractions from size-exclusion chromatography. The bands corresponding to NALCN and FAM155A protein were indicated. The fractions indicated with red line were used for cryo-EM sample preparation.

**Fig. S2.**
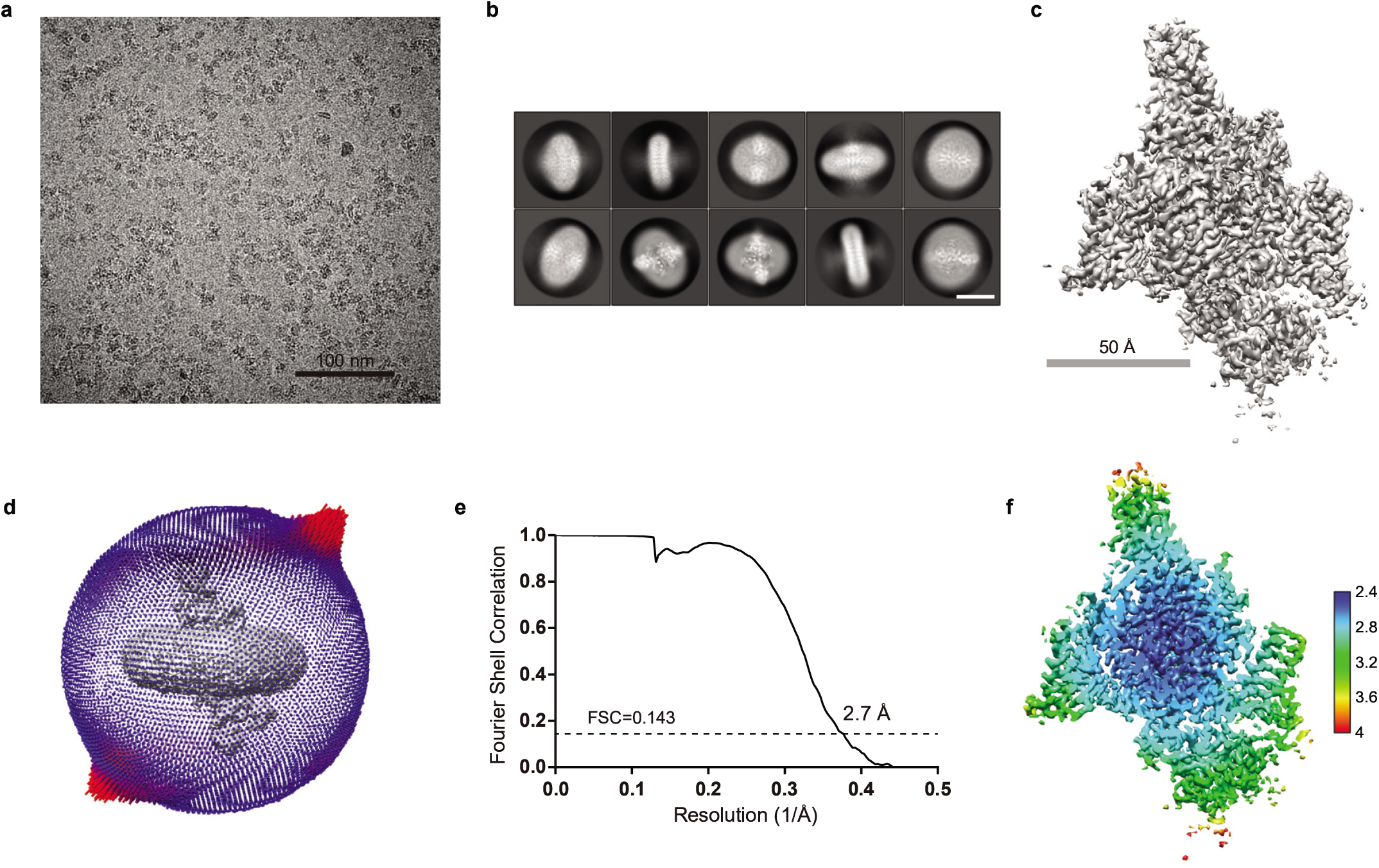
Cryo-EM analysis of NALCN-FAM155A complex. **a**, Representative raw micrograph of NALCN-FAM155A complex. **b**, Representative 2D class average of NALCN-FAM155A particles. Scale bar, 10 nm. **c**, Cryo-EM map of NALCN-FAM155A complex. **d**, The angular distribution of final reconstruction. **e**, Gold standard FSC curve of NALCN-FAM155A EM map. Resolution estimations are based on the criterion of FSC 0.143 cutoff. **f**, Local resolution map of NALCN-FAM155A complex calculated using Relion 3.1.

**Fig. S3.**
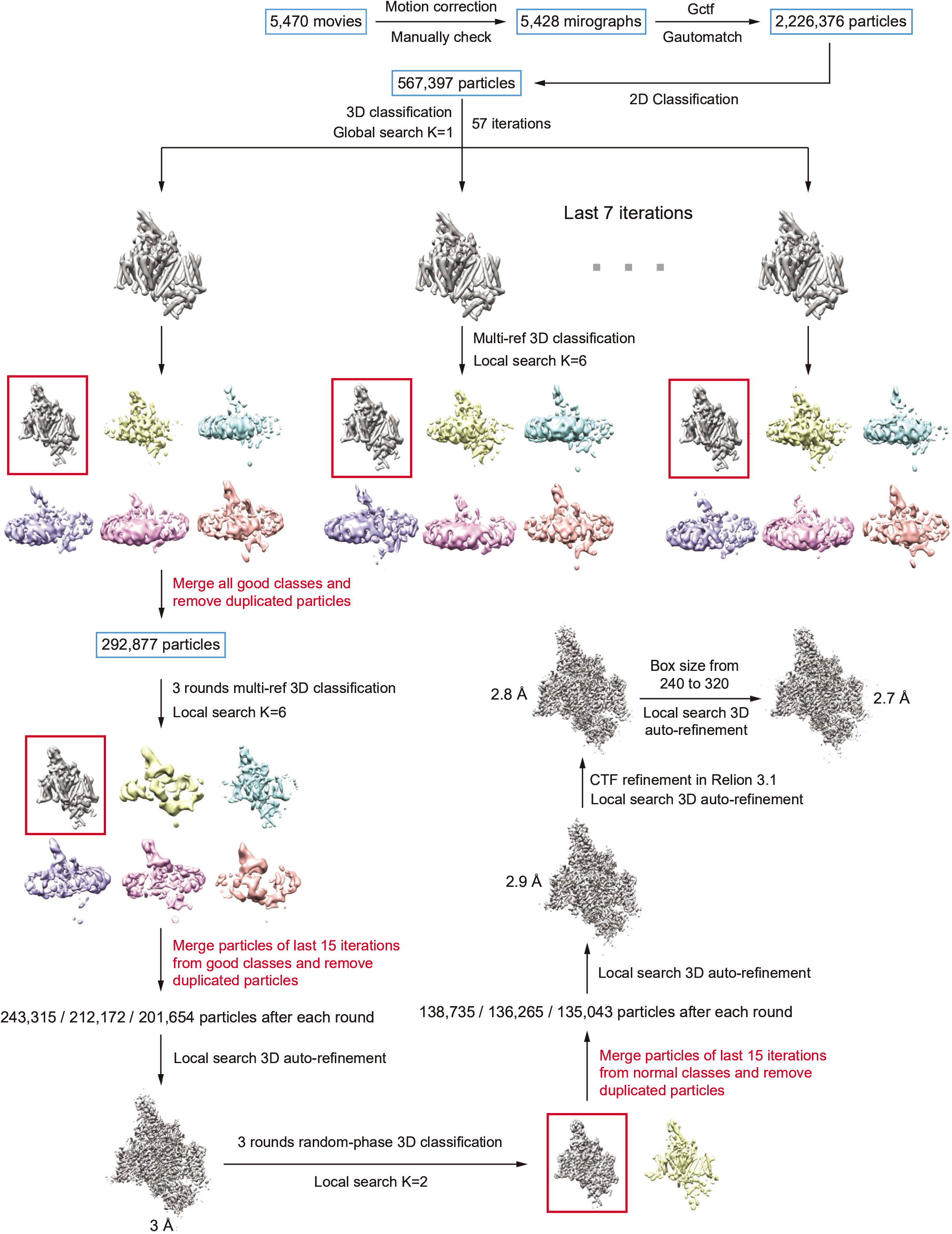
Workflow for cryo-EM data processing of NALCN-FAM155A complex. For details, see ‘Image processing’ in Methods section.

**Fig. S4.**
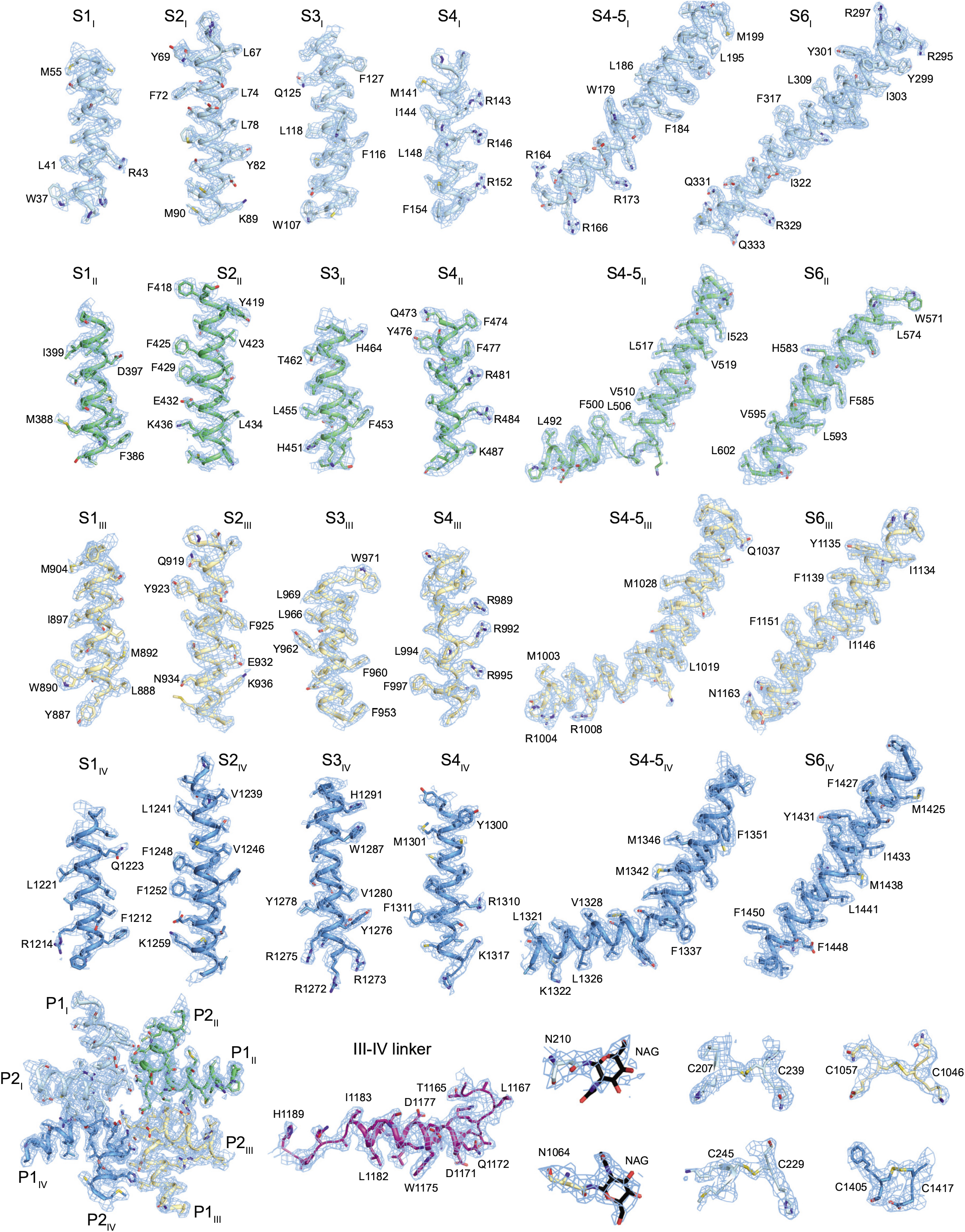
Representative EM maps for segments in NALCN subunit. EM maps for S1-S6 transmembrane segments, pore domain, III-IV linker, sugar moiety and disulfide bonds in NALCN subunit, shown as blue mesh, are contoured at 3.5 σ.

**Fig. S5.**
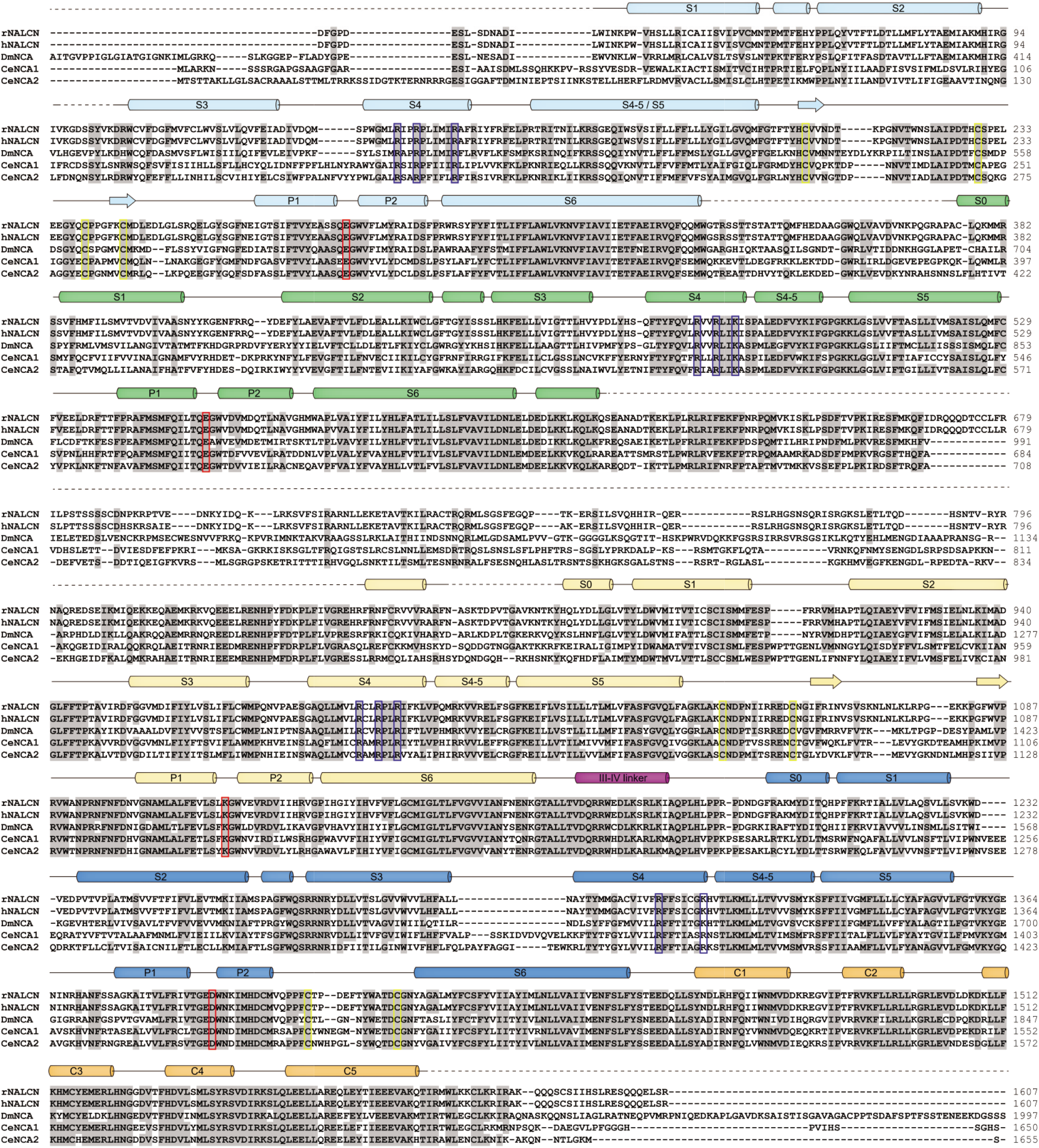
Sequence alignment of NALCN subunit. The sequences of rat NALCN, human NALCN, fruit fly NCA, nematode NCA1 and NCA2 were aligned. Conserved residues are shaded in grey. Residues forming disulfide bonds, the selectivity filter “EEKD”, and the positive residues on S4 segments are indicated by yellow, red, and blue boxes, respectively. Secondary structural elements are indicated as follows: arrows, β-sheets; cylinders, α-helices; lines, loops. Disordered regions are shown as dashed lines.

**Fig. S6.**
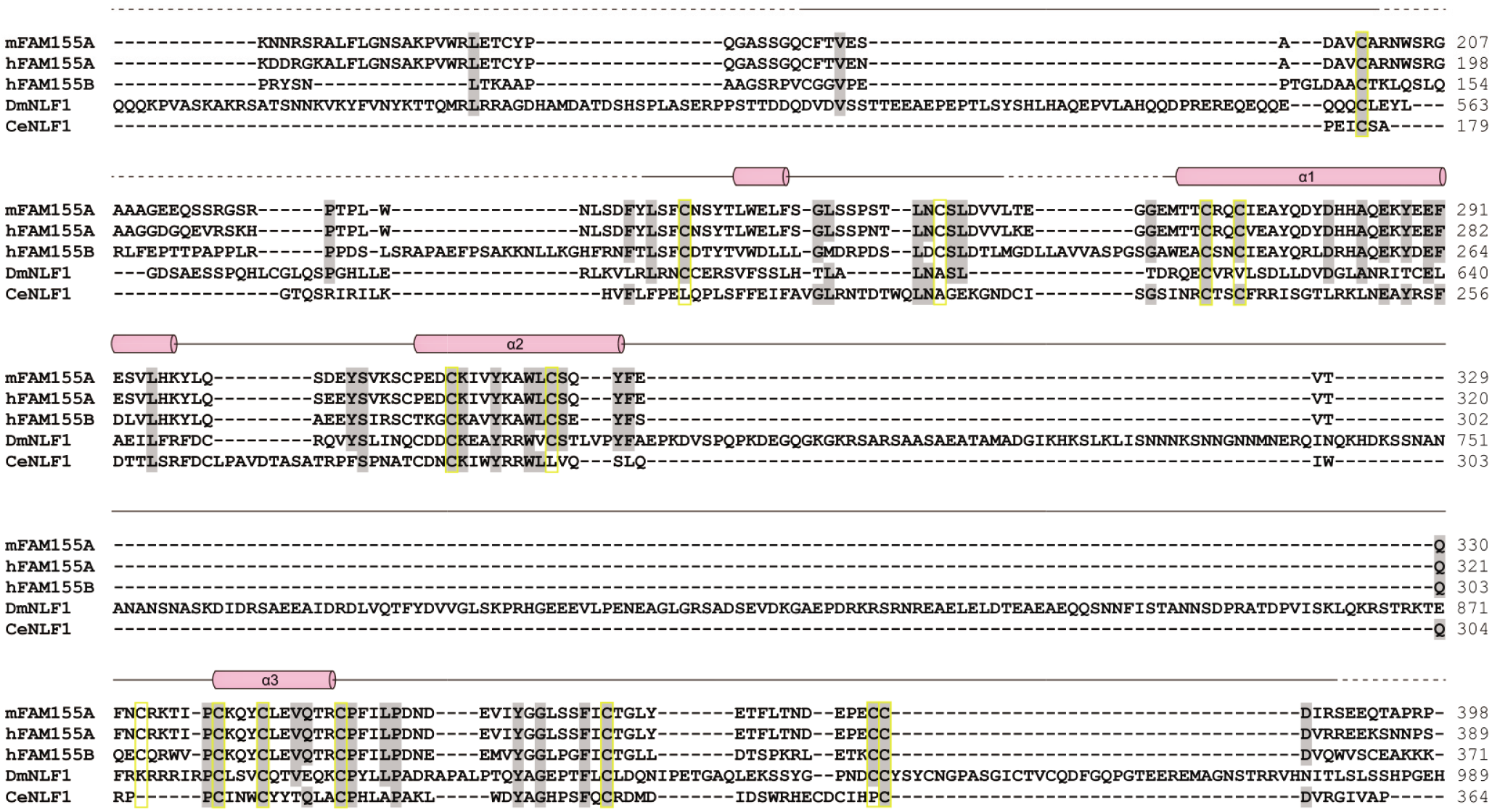
Sequence alignment of FAM155A subunit. The sequences of mouse FAM155A, human FAM155A, human FAM155B, fruit fly NLF1 and nematode NLF1 were aligned. Conserved residues are shaded in grey. Residues forming disulfide bonds are indicated by yellow boxes. Secondary structural elements are indicated as follows: cylinders, α-helices; lines, loops. Disordered regions are shown as dashed lines.

**Fig. S7.**
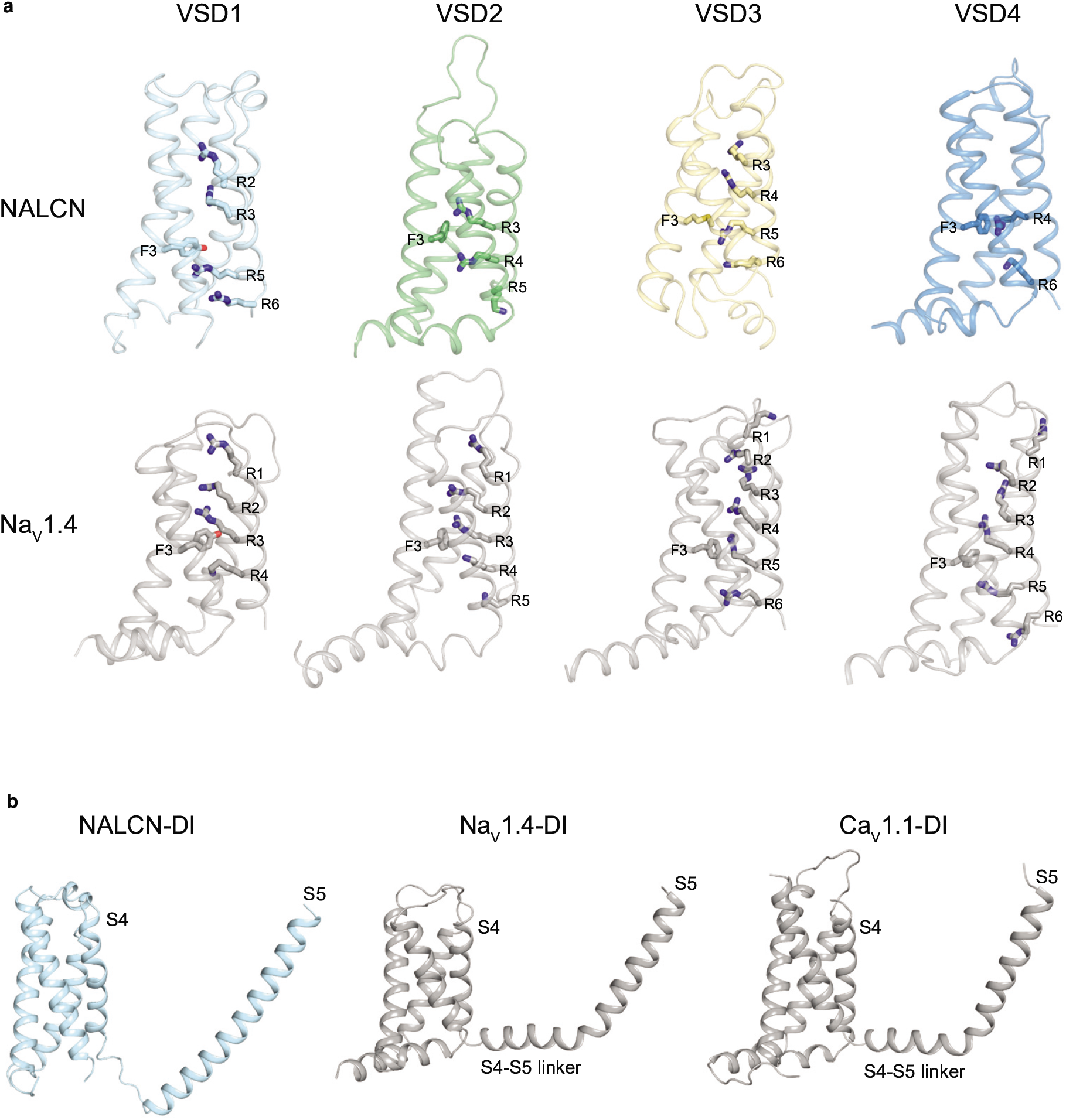
NALCN voltage-sensor domains and S4-5 linker comparisons. **a**, Comparison of the VSDs from NALCN and Na_V_1.4 (PDB: 6AGF). **b**, Comparison of the DI-S4-5 linker from NALCN, Na_V_1.4 (PDB: 6AGF) and Ca_V_1.1 (PDB: 5GJV). Only S0-S5 segments of DI were shown for clarity.

**Fig. S8.**
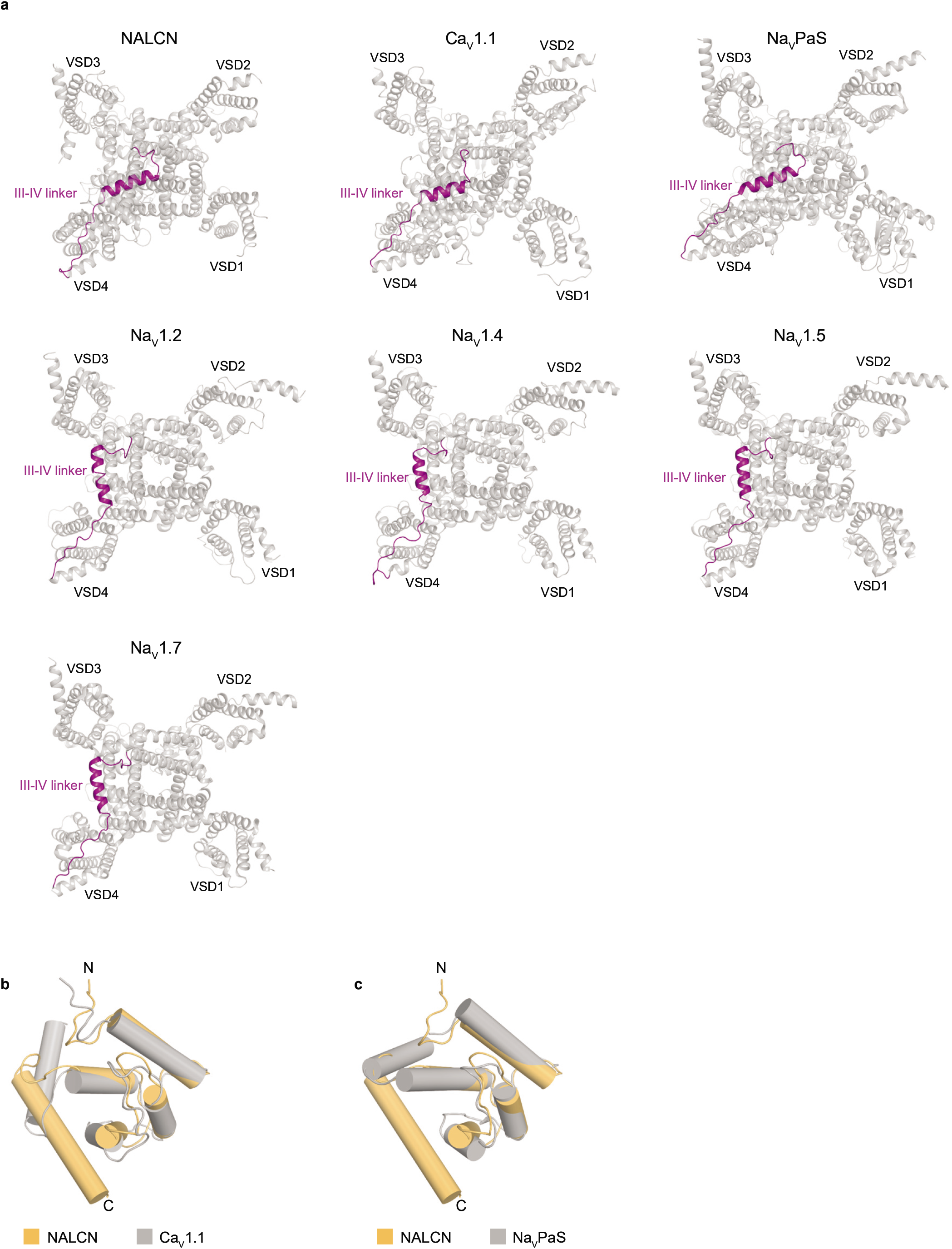
NALCN III-IV linker and C-terminal domain comparisons. **a**, Comparison of III-IV linker (bottom view) from NALCN, Ca_V_1.1 (PDB: 5GJV), Na_V_PaS (PDB: 5X0M), Na_V_1.2 (PDB: 6J8E), Na_V_1.4 (PDB: 6AGF), Na_V_1.5 (PDB: 6UZ3) and Na_V_1.7 (PDB: 6J8G). III-IV linkers are colored by purple. **b**, Comparison of the CTD from NALCN, Ca_V_1.1 (PDB: 5GJV) and Na_V_PaS (PDB: 5X0M). Helices are shown as cylinders.

## Acknowledgements

We thank Prof. Dejian Ren for providing the cDNA of hNALCN, rNALCN, mUNC80, and mUNC79, Dr. Lear Bridget for providing the cDNA of DmUNC79 and DmUNC80, Dr. Ravi Allada for providing the cDNA of DmNCA and DmNLF-1, Dr. Yuji Kohara for providing the cDNA of CeNCA1. Cryo-EM data collection was supported by Electron microscopy laboratory and Cryo-EM platform of Peking University with the assistance of Xuemei Li, Daqi Yu, Xia Pei, Bo Shao, Guopeng Wang, and Zhenxi Guo. Part of structural computation was also performed on the Computing Platform of the Center for Life Science and High-performance Computing Platform of Peking University. This work is supported by grants from the Ministry of Science and Technology of China (National Key R&D Program of China, 2016YFA0502004 to L.C.), National Natural Science Foundation of China (91957201, 31622021, 31870833 and 31821091 to L.C.), Beijing Natural Science Foundation (5192009 to L.C.), and Young Thousand Talents Program of China to L.C.

## Author contributions

Lei Chen initiated the project. Yunlu Kang did the electrophysiology recording, purified proteins and prepared the cryo-EM samples. Yunlu Kang and Jing-Xiang Wu collected the cryo-EM data. Yunlu Kang processed the cryo-EM data. Yunlu Kang and Lei Chen built and refined the atomic model. All authors contributed to the manuscript preparation.

---

**Extended Data Table 1.** Cryo-EM data collection, refinement and validation statistics

|  |  |
| --- | --- |
|  | NALCN-FAM155A |
| PDB ID | XXX |
| EMDB ID | XXX |
| **Data collection and processing** |  |
| Magnification | 130,000 × |
| Voltage (kV) | 300 |
| Electron exposure (e<sup>-</sup>/Å<sup>2</sup>) | 50.9 |
| Defocus range (μm) | -1.5 to -1.8 |
| Pixel size (Å) | 1.045 |
| Symmetry imposed | *C1* |
| Final particle images (no.) | 135,043 |
| Map resolution (Å) | 2.7 |
| FSC threshold | 0.143 |
| Map sharpening *B* factor (Å<sup>2</sup>) | -78.6 |
| **Refinement** |  |
| Model composition |  |
| Non-hydrogen atoms | 12,181 |
| *B* factors (Å<sup>2</sup>) | 45.35 |
| R.m.s. deviations |  |
| Bond lengths (Å) | 0.006 |
| Bond angles (°) | 0.629 |
| Validation |  |
| MolProbity score | 2.07 |
| Clashscore | 6.22 |
| Poor rotamers (%) | 3.26 |
| Ramachandran plot |  |
| Favored (%) | 95.20 |
| Allowed (%) | 4.80 |
| Disallowed (%) | 0.00 |

